# The role of fucose-containing glycan motifs across taxonomic kingdoms

**DOI:** 10.1101/2021.08.08.455599

**Authors:** Luc Thomès, Daniel Bojar

## Abstract

The extraordinary diversity of glycans leads to large differences in the glycomes of different kingdoms of life. Yet, while most monosaccharides are solely found in certain taxonomic groups, there is a small set of monosaccharides with widespread distribution across nearly all domains of life. These general monosaccharides are particularly relevant for glycan motifs, as they can readily be used by commensals and pathogens to mimic host glycans or hijack existing glycan recognition systems. Among these, the monosaccharide fucose is especially interesting, as it frequently presents itself as a terminal monosaccharide, primed for interaction with proteins. Here, we analyze fucose-containing glycan motifs across all taxonomic kingdoms. Using a hereby presented large species-specific glycan dataset and a plethora of methods for glycan-focused bioinformatics and machine learning, we identify characteristic as well as shared fucose-containing glycan motifs for various taxonomic groups, demonstrating clear differences in fucose usage. Even within domains, fucose is used differentially based on an organism’s physiology and habitat. We particularly highlight differences in fucose-containing motifs between vertebrates and invertebrates. With the example of pathogenic and non-pathogenic *Escherichia coli* strains, we also demonstrate the importance of fucose-containing motifs in molecular mimicry and thereby pathogenic potential. Finally, we also confirm and extend the role of fucosyltransferase-coding genes (FUT) in several important biological processes that include development, immunity, and diseases, especially cancer. We envision that this study will shed light on an important class of glycan motifs, with potential new insights into the role of fucosylated glycans in symbiosis, pathogenicity, and immunity.

## INTRODUCTION

Biological polysaccharides (or glycans) were long considered as solely nutrition and energy supply. Since decades, however, it has been shown that they can be covalently linked to proteins, through the important post-translational mechanism known as glycosylation. Despite their well-characterized association with proteins, their functional roles often remained elusive. Recently, thanks to constant progress in glycobiology, more insights were gained about the impact of glycan architecture and composition on their functions in various biological processes. On proteins, they are important to attain and maintain their folded structure (Shental-Bechor and Levy, 2008). At the cell level, glycosylated proteins are targeted to the membrane or secreted into the extracellular matrix where they, for instance, form a physical barrier that can store secreted factors, modulating their accessibility and establishing interactions with proteins or cells (Lindahl et al., 2017). Glycans are also important for mediating inter-organism communication, including the initiation of viral contact with host cells (Watanabe et al., 2020). They are especially involved in immune response regulation, as they can be targeted by glycan-binding immune proteins and modulate the efficacy of antibodies (Li et al., 2021).

Additionally, glycans are found on lipids (Hove et al., 2021) and, as recently detailed, even on cell-surface RNAs (Flynn et al., 2021). The omnipresence of polysaccharides on fundamental building blocks demonstrates their necessity across all levels of biological systems. This involvement in such diverse biological functions is facilitated by i) the high combinatorial potential of chained monosaccharides resulting in a large diversity of glycans with specific properties, ii) their presence on all kinds of cellular components, cell types, and tissues, and iii) their conservation during evolution that resulted in their ubiquitous presence across all taxonomic kingdoms. In contrast to the wide distribution of glycans, most of their constitutive monosaccharides are solely found in specific taxonomic groups (Srivastava et al., 2020). For example, sialic acids (often found in glycans as N-acetyl-5-neuraminic acid or Neu5Ac) are prominent in vertebrates but usually considered absent from most invertebrates, plants, and prokaryotic organisms (Angata and Varki, 2002; Schauer, 2004). Only a rather small set of these fundamental building blocks have a widespread distribution across nearly all domains of life. Fucose (Fuc) is one of these almost-universal monosaccharides, making it particularly suited to serve as a common language element between different organisms.

Contrary to the standard D-hexoses commonly found in glycans, such as glucose or mannose, fucose is a deoxyhexose that is mostly present in the L configuration. Often a terminal monosaccharide in polysaccharides, it is particularly well suited to mediate protein interactions which are essential to many biological processes including host-microbiota communication, viral infection, or immunity. In mammals, its importance is fundamental. Fucosylation constitutes a component of the ABO blood group and the Lewis antigen systems. It consists of transferring a fucose residue from a guanosine diphosphate (GDP)-fucose donor to an acceptor molecule. Depending on the site of fucose addition catalyzed by the fucosyltransferases (FUTs), fucosylations are classified into α1-2-, α1-3/4-, α1-6-, and *O*-fucosylation (Schneider et al., 2017). In humans, these reactions are performed by a set of thirteen enzymes, each displaying a specific activity. From these, FUT2 and FUT3 play major roles in the Lewis antigen system, as they are responsible for the production of Le^a^, Type 1 H, and Le^b^ antigens secreted in body fluids, including plasma. These Le^b^ antigens are of notable importance, as their expression at the surface of epithelial cells in the stomach is recognized and exploited by the bacterium *Helicobacter pylori*, to adhere to the epithelium and escape the surrounding hostile acidic environment (Bugaytsova et al., 2017; Matos et al., 2021).

In animals, FUT8 is widely studied and is responsible for the formation of the widespread core fucose motif: fucose connected to the core chitobiose of *N*-linked glycans at the reducing end through an α1-6 linkage. Many studies based on genetic engineering or fucosylation inhibitors have highlighted the importance of this motif in animals. As FUT8 encodes the only animal enzyme capable of adding fucose to GlcNAc residues via an α1-6 linkage, studies demonstrated that its deletion by knock-out (FUT8^−/−^) in mice results in major impairments, leading to growth retardation and premature death (Wang et al. 2005). Similarly, in humans, mutations leading to an abnormal or complete loss of function of the encoded enzyme result in a congenital disorder of glycosylation (CDGF1) associated with similar symptoms, including a general developmental delay (Ng et al. 2018). The human FUT8 gene is susceptible to single nucleotide polymorphisms (SNPs) that have an impact on the amount of core fucosylation in maternal milk *N*-glycans. A decreased level of these modifications in milk results in alterations of breast-fed infants’ microbiota (Li et al., 2019), favoring the development of allergies and metabolic diseases (Tamburini et al., 2016). In immunoglobulins, core fucosylation is particularly important. IgG antibodies direct eosinophils and natural killer cells towards cancer and pathogen-infected cells. This process, referred to as antibody-dependent cell-mediated cytotoxicity (ADCC), involves the binding of antibodies to effector and target cells by the constant (Fc) and variable regions of the antibody, respectively (Li et al., 2021). This activity is influenced by glycans present on the Fc region of IgG that affect its function and folding, with a lack of fucosylation being associated with an increased capacity to connect the antibody constant region to the antibody receptor FcγRIIIa.

Based on the conserved nature of glycan-mediated interspecies communication (Wanke et al., 2021), we decided to study fucose and its context in glycans across all taxonomic groups using the glycowork package (Thomès et al., 2021). Because of the complexity of glycans, manual investigations are struggling to extract meaning from these constituents. To solve this problem, an increasing number of *in silico* approaches dedicated to their analysis have been recently developed, including molecular dynamics simulations of protein-associated glycans (Harbison et al., 2019; Fogarty et al., 2020) or glycan-focused machine learning efforts to link sequences to functions (Bojar et al., 2020b; Burkholz et al., 2021). Bioinformatics methods are well-suited to investigate such complex molecules in an efficient and automated manner, particularly with the synergy between classical computational approaches with pattern-finding machine learning models. Here, we investigated the importance of fucosylation and the associated enzymes for biological processes, including symbiosis and infection. We demonstrated that fucose-containing glycans are present in all kingdoms and that the ratio between fucose-containing and total glycans within a kingdom or a species is often informative. One example of this is found in bacteria, where we correlated this ratio with the capacity of organisms to grow in different environments, to display pathogenic activity, and to evade the host immune system through a mimicking process. We also analyzed the relation of animal fucose-related enzymes with major diseases including cancer. Overall, our analyses illuminate a panoply of properties and functions across taxonomic kingdoms that rely on fucose-containing motifs, emphasizing the importance of this monosaccharide.

## MATERIALS AND METHODS

### Dataset

The data used in this study were based on a previously reported dataset (Bojar et al., 2020a; Bojar et al., 2021) that we updated for this work by the aggregation of data from public databases (GlyTouCan (Fujita et al., 2021), GlyCosmos (Watanabe et al., 2021), CSDB (Toukach and Egorova, 2016)), together with glycan structures manually extracted from the peer-reviewed literature. The updated dataset presented and analyzed here contained a total of 22,888 glycan sequences in the IUPAC-condensed nomenclature, associated with the lineage information of the 2,171 species from which they stemmed. This dataset is released as a part of the glycowork 0.2 package, an updated version of the work from (Thomès et al., 2021), and is freely accessible via https://github.com/BojarLab/glycowork.

### Machine learning model training to obtain learned similarities

To use the motif.analysis.plot_embeddings() function of glycowork, we trained SweetNet-type machine learning models (Burkholz et al., 2021) to obtain glycan representations. Two models were trained for this. One based on all the glycans from the dataset presented here and a second after including *Escherichia coli* glycans associated with their pathogenic potential, enabling comparisons between pathogenic, non-pathogenic, and uncharacterized strains. For both models, we randomly split our data into 80/20% for train and test sets, respectively. Glycan representations or learned similarities were obtained after the graph convolutional layers of the trained neural network, as described in Burkholz et al., 2021.

### Data Pre-processing

To improve the readability of our figures, the different kingdoms initially found in our dataset were simplified as follows. The “Virus”, “Orthornavirae”, “Riboviria”, and “Heunggongvirae” kingdoms were merged into a unique “Virus” group. Similarly, “Protista”, “Excavata”, and “Chromista” were grouped under the “Protista” kingdom, and “Crenarchaeota”, “Euryarchaeota”, and “Proteoarchaeota” were merged into an “Archaea” group.

### Visualizing glycan properties via embeddings and heatmaps

All data were analyzed using the functions implemented in Glycowork. T-SNE graphs (van der Maaten and Hinton, 2008) were generated with the motif.analysis.plot_embeddings() function according to the glycan representation obtained during the machine learning training step. Motif-based heatmaps were computed using the motif.analysis.make_heatmap() function of glycowork.

### Fucose usage across bacterial species

To make the computation of the proportion of fucosylated glycans across different bacterial species more robust, we applied a filter based on the number of available glycan structures. Bacteria with too low number of glycans in our database can reach extremely high or low fucosylated glycans proportions that are not biologically representative but rather reflect our current knowledge. Consequently, we only kept species with at least 30 characterized structures. For these species, information about preferred growing environment and disease association were retrieved from the peer-reviewed literature.

### Pathogenic potential of *Escherichia coli* strains

For the study of *E. coli* pathogenic potential, we generated glycan representations using the motif.analysis.plot_embeddings() function, based on a previously reported dataset (Bojar et al., 2021) flagging each strain with a value of 0 (non-pathogenic) or 1 (pathogenic). To characterize statistically enriched motifs from fucose-containing glycans of pathogenic versus non-pathogenic and non-pathogenic versus pathogenic strains, we applied the motif.analysis.get_pvals_motifs() function from glycowork, with a threshold of 0, which calculates motif enrichment based on one-tailed Welch’s t-tests with a Holm-Šídák correction. Only motifs with p-values below 0.05 were considered.

### Data mining using the EMBL-EBI Expression Atlas

To interrogate the EMBL-EBI Expression Atlas website in an automated fashion, we used an in-house Python script taking a gene list as input and sending requests to the database. For each gene encoding a human fucosyltransferase (FUT1 to FUT11, POFUT1, and POFUT2), the script extracted all dataset names, compared conditions, log2 fold-changes, and adjusted p-values in which the concerned gene is differentially expressed. At the end of this step, a final table is produced, listing how many and which of the thirteen human fucosyltransferases genes are differentially expressed for all experiments. Only genes with an adjusted p-value inferior to 0.05, a log2 fold-change superior or equal to 1 or, otherwise, inferior or equal to −1 were considered differentially expressed. We focused this analysis on animal genes and consequently discarded homonymous genes from non-animal species.

## RESULTS

### Investigation of fucose distribution in glycans across taxonomic kingdoms

Fucose is a monosaccharide that is considered widespread across all taxonomic kingdoms. In our new, comprehensive dataset of 22,888 species-specific glycans, we found fucose-containing glycans among all seven investigated kingdoms, Animalia, Plantae, Fungi, Bacteria, Protista, Virus, and Archaea (Fig. 1A). This observation was consistent with its known status as a widespread monosaccharide. We quantified the proportion of fucosylated versus total glycans among these groups and demonstrated that the highest proportions of fucose-containing glycans were observed in animals and viruses, while the lowest proportions were observed in Fungi, Protista, and Archaea. An intermediate situation was observed in Plantae and Bacteria (Fig. 1B). Interestingly, the only protist that contained fucosylated glycans in our dataset was *Dictyostelium discoida*, the unique representant of the amoebozoa phylum studied here. Similarly, *Thermoplasma acidophila* was the only archaeal species from our dataset that exhibited fucosylated glycans. Based on these observations, further efforts in characterizing these and other taxonomic groups are needed to decipher whether these species constitute exceptions among their groups or whether fucosylation is characteristic of their respective underrepresented phylum and order.

**Figure 1.**
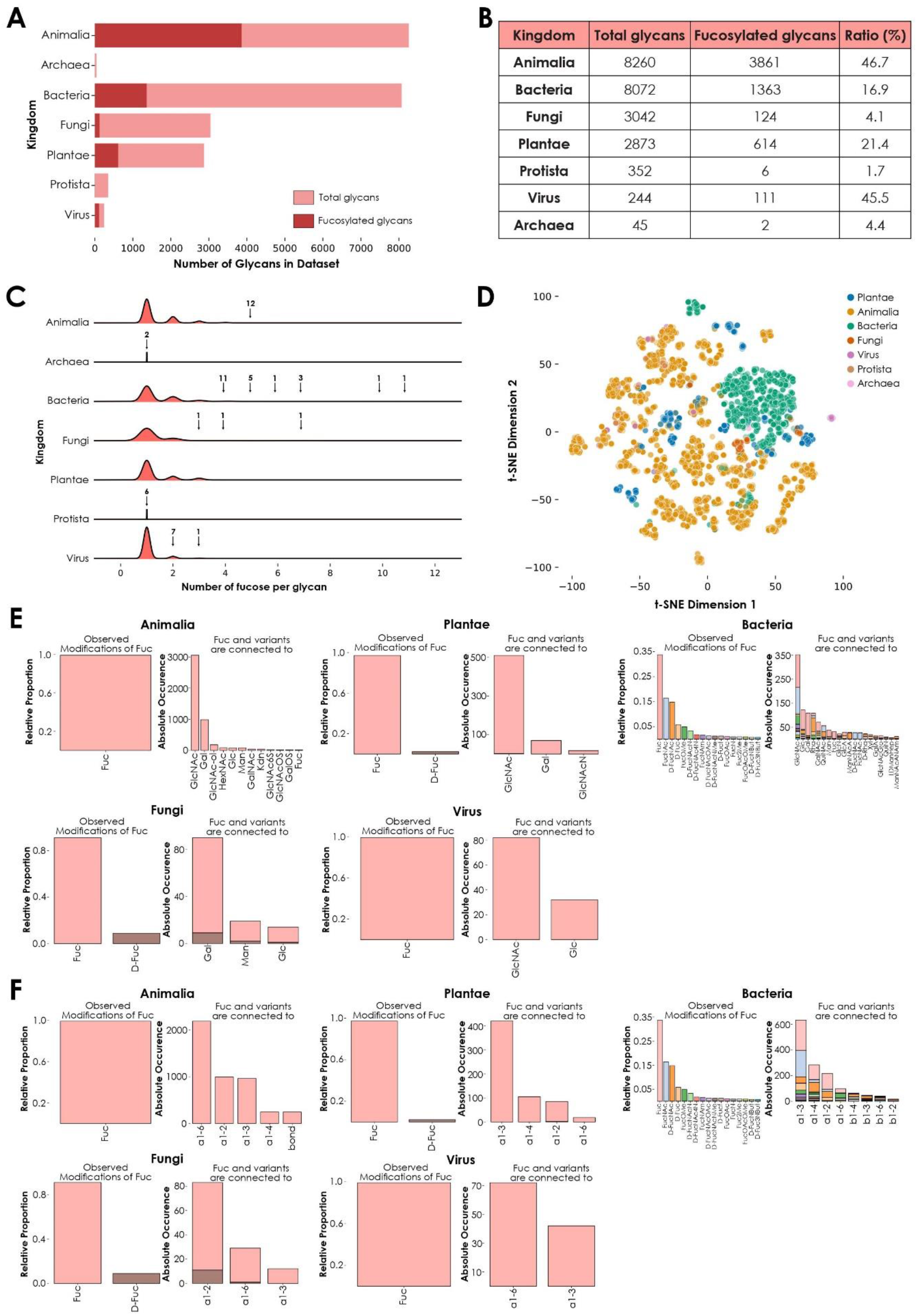
Kingdom-level descriptions of fucose usage. **A)** Number of fucose-containing glycans across kingdoms. Total number of glycans available in the dataset for each kingdom are represented in light red. The total number of fucose-containing glycans is depicted in dark red. **B)** Ratio of total versus fucosylated glycans. Number of total and fucosylated glycans from the seven investigated kingdoms are ordered by decreasing number of total glycans. The ratio of fucosylated glycans is indicated as a percentage. **C)** Fucose enrichment in glycans. This joyplot shows, for each kingdom, the number of fucose units per fucose-containing glycan. Arrows indicate peaks corresponding to fucosylated glycans present in too low proportions to be visible and their associated number of glycans. **D)** Diversity of fucose-containing glycans across kingdoms. Fucose-containing glycans from every species in the dataset are represented as colored dots via t-SNE, based on their learned similarities from a SweetNet model. Colors correspond to the taxonomic kingdom from which they stem. **E)** Usage of fucose variants and preferred neighbors. Each graph represents the relative abundance and characteristics of fucose and fucose variants. Colored stacked bars correspond to the preferred neighbors of fucose variants in glycans, with colors corresponding to the variants observed on the left part of each graph. **F)** Preferred linkage-type between fucose and other monosaccharides. Each graph represents the relative proportion of linkage-types that connect fucose and fucose variants with other monosaccharides in glycans. Colored stacked bars correspond to the linkage preferences of fucose variants in glycans, with colors corresponding to the variants.

To further investigate this heterogeneous distribution across kingdoms, we then counted the occurrence of fucose residues within each glycan from these groups (Fig. 1C). Intriguingly, glycans with the highest fucosylation were found in bacterial (up to eleven fucose units) and fungal (up to seven fucose units) species. In animals, where the proportion of fucose-containing glycans is the highest (Fig. 1B), the maximal number of fucoses per glycan was five. This observation indicated that the number of fucosylated glycans found within a kingdom did not necessarily correlate with the maximal number of fucose units in glycans observed from this kingdom.

We also analyzed the kingdom-specific properties of fucosylated glycans, especially their composition and structure. We hypothesized that these two parameters were responsible for the large variance of fucose-containing glycans seen on a t-SNE plot using learned glycan representations (Fig. 1D). Fucosylated glycans found within each kingdom share a particular degree of similarity, as indicated by their respective clustering. This observation suggested the existence of common biosynthetic pathways. Kingdom-specific pathways are known to involve enzymes responsible for the integration of fucose monosaccharides and fucose variants in different glycan contexts (Fig. 1E) and via specific linkages (Fig. 1F) (Oriol et al., 1999; Roos et al., 2002). Strikingly, fucose was always preferentially connected to N-acetyl glucosamine (GlcNAc), except for the Fungi kingdom where it was more often linked to galactose (Gal) instead. Also, while most kingdoms use L-fucose or, to a lesser extent, D-fucose, bacteria displayed a many fucose variants, including L-FucNAc (N-acetyl fucosamine) and D-FucNAc, in a high proportion. In terms of linkage-type, each kingdom exhibited its own specificities. In plants and bacteria, the most frequent linkage between fucose and its neighbors was the α1-3 bond. In animals and fungi, the most preferred linkages were α1-6 and α1-2, respectively. Viruses used both α1-6 and α1-3 linkages in high proportions, mirroring the preferences observed in their respective animal and plant hosts as, except for a few virus families that perform autonomous glycosylation (Piacente et al., 2015), all known viral particles are decorated with glycans synthetized by the host cell machinery.

### Fucose-containing glycans from invertebrate species

We hypothesized that discernible trends in fucose usage also differed in taxonomic groups below the kingdom level. We chose to focus on the Animalia kingdom to see whether fucosylated glycans were sufficient to understand intra-kingdom variability. One major dividing line within this kingdom emerges from comparing vertebrates and invertebrates. For example, it is well known that sialic acid is rare in many invertebrates which, consequently, produce less sialylated glycans (for a review, see Paschinger and Wilson, 2019).

Our analyses yielded that the difference between these two groups of organisms was also reflected in their usage of fucosylated glycans (Fig. 2). The representation of glycans from vertebrates (all found within the Chordata phylum) and invertebrates (all other animal phyla) showed that glycans from the latter clearly formed separate clusters (Fig. 2A). As the position of glycans on this plot is essentially due to their composition and architecture, we investigated these features on a heatmap representing the most abundant monosaccharides, linkages, and motifs observed within fucosylated glycans (Fig. 2B). Many motifs were found exclusively in vertebrates, mirroring the usage of sialylated glycans in these organisms. Some of the most striking differences concerned internal and terminal LacNAc (N-acetyllactosamine) type 2 motifs, comprising a Gal(ß1-4)GlcNAc disaccharide, that predominated in vertebrates. In addition, the heatmap also confirmed the specific presence of Neu5Ac as a characteristic feature of Chordata, as described in the literature (Schauer, 2004).

**Figure 2.**
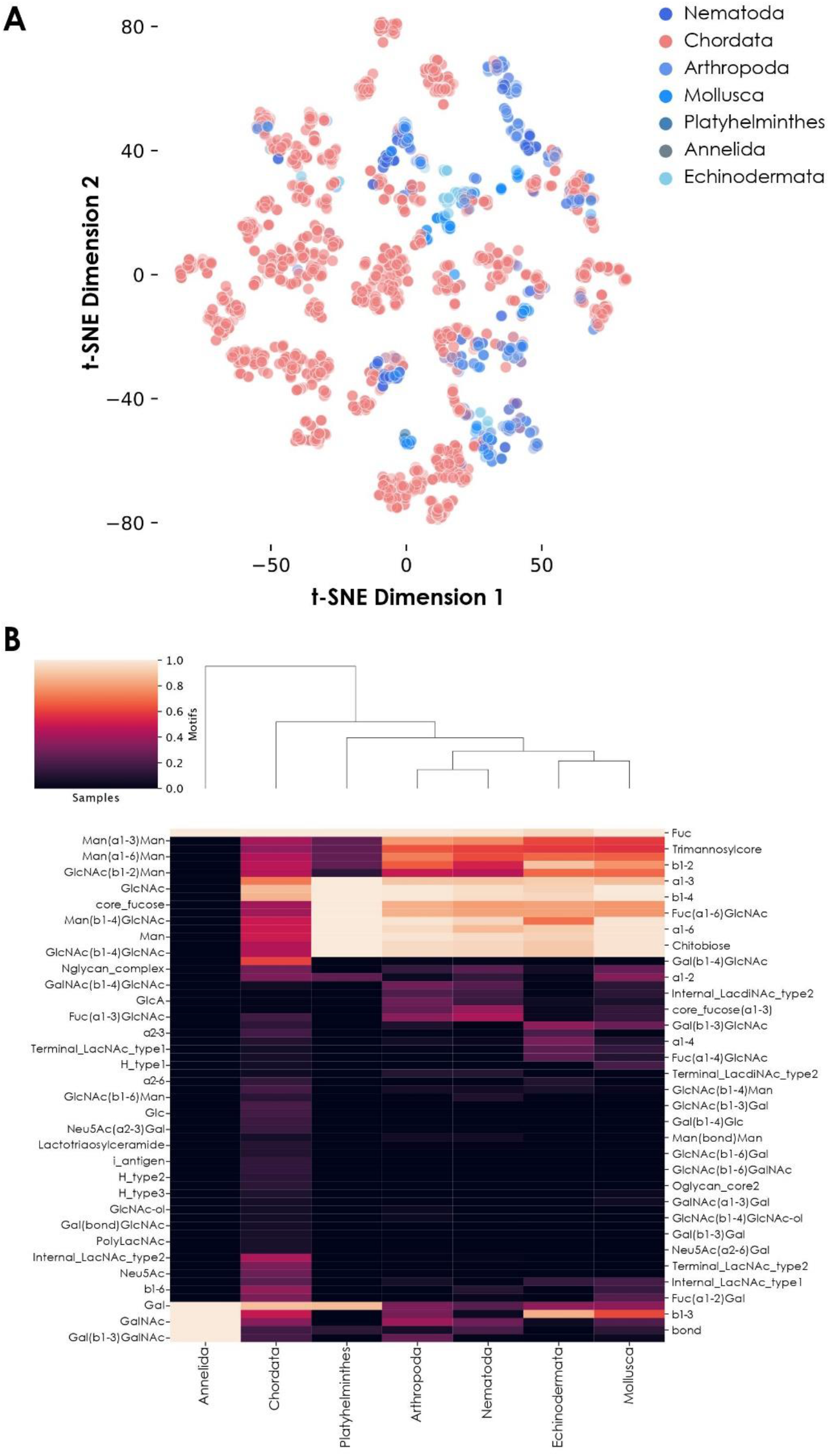
Fucose usage in animal glycans. **A)** Diversity of fucose-containing glycans among animals. Fucose-containing glycans from animal species are represented via t-SNE as colored dots, based on their learned similarity from a SweetNet model. Colors indicate the phylum from which they stem. **B)** Motif distribution from fucosylated glycans across animal phyla. This heatmap represents the relative abundance of different glycan motifs in each of the seven studied animal groups. Notably, Annelida only contains one fucosylated glycan in our dataset.

Interestingly, some motifs and monosaccharides were more characteristic of invertebrate fucosylated glycans. This included internal LacdiNAc type 2 (N,N-Diacetyllactosamine; GalNAc(ß1-4)GlcNAc) and core fucose (α1-3) motifs, as well as the presence of glucuronic acid (GlcA) (Fig. 2B). These observations were consistent with previous detection of GlcA- and LacdiNAc-containing glycans from different insect species (Kurz et al., 2015). Also, the presence of mammalian-like α1-6 and plant-like α1-3 core fucose motifs in these organisms, isolated and in combination, is another specificity initially observed in honeybee venom (Kubelka et al., 1993). While Chordata glycans lack the core fucose (α1-3) motif, they do contain the Fuc(α1-3)GlcNAc disaccharide motif on *N-* and *O*-linked glycans. This observation suggests an alternative usage of Fuc(α1-3) in vertebrates which, contrary to invertebrates, do not use it to decorate the reducing monosaccharide at the base of *N*-glycans. Fucose-containing glycans from the Annelida phylum seemed to display a highly characteristic profile, in which many features shared between Chordata and invertebrates were absent, including any of the core fucose types classically found in animals. However, a closer inspection of the data revealed that only one glycan from the four Annelida species in our dataset is fucosylated, preventing any conclusions.

### Fucosylation of *N*-linked and *O*-linked glycans from animals

*N*-linked and *O*-linked glycans are the most frequent types of polysaccharides found linked to animal proteins via post-translational glycosylation. To further analyze observed differences between vertebrates and invertebrates, we separated all their fucosylated glycans into *N*- and *O*-linked glycans based on the presence of GlcNAc or GalNAc at the reducing end, respectively (Fig. 3). We noted the presence of many more *N-*linked (Fig. 3A) than *O-*linked (Fig. 3B) fucosylated glycans in invertebrates compared to vertebrates. To quantify this depletion of fucosylated *O*-linked glycans in invertebrates, we computed the proportion of *O-*linked fucosylated versus total *O*-linked glycans in vertebrates and invertebrates (Fig. 3C). This analysis showed that half of the vertebrate *O-*glycans are fucosylated while, in comparison, only 27.9% of the invertebrate *O*-glycans are. While we found almost 22 times more *O*-linked glycans in vertebrates than in invertebrates in our dataset, there are 39 times more *O*-linked fucosylated glycans in the former group compared to the latter. These results clearly indicated a substantial enrichment of fucosylated glycans in vertebrates compared to invertebrates.

**Figure 3.**
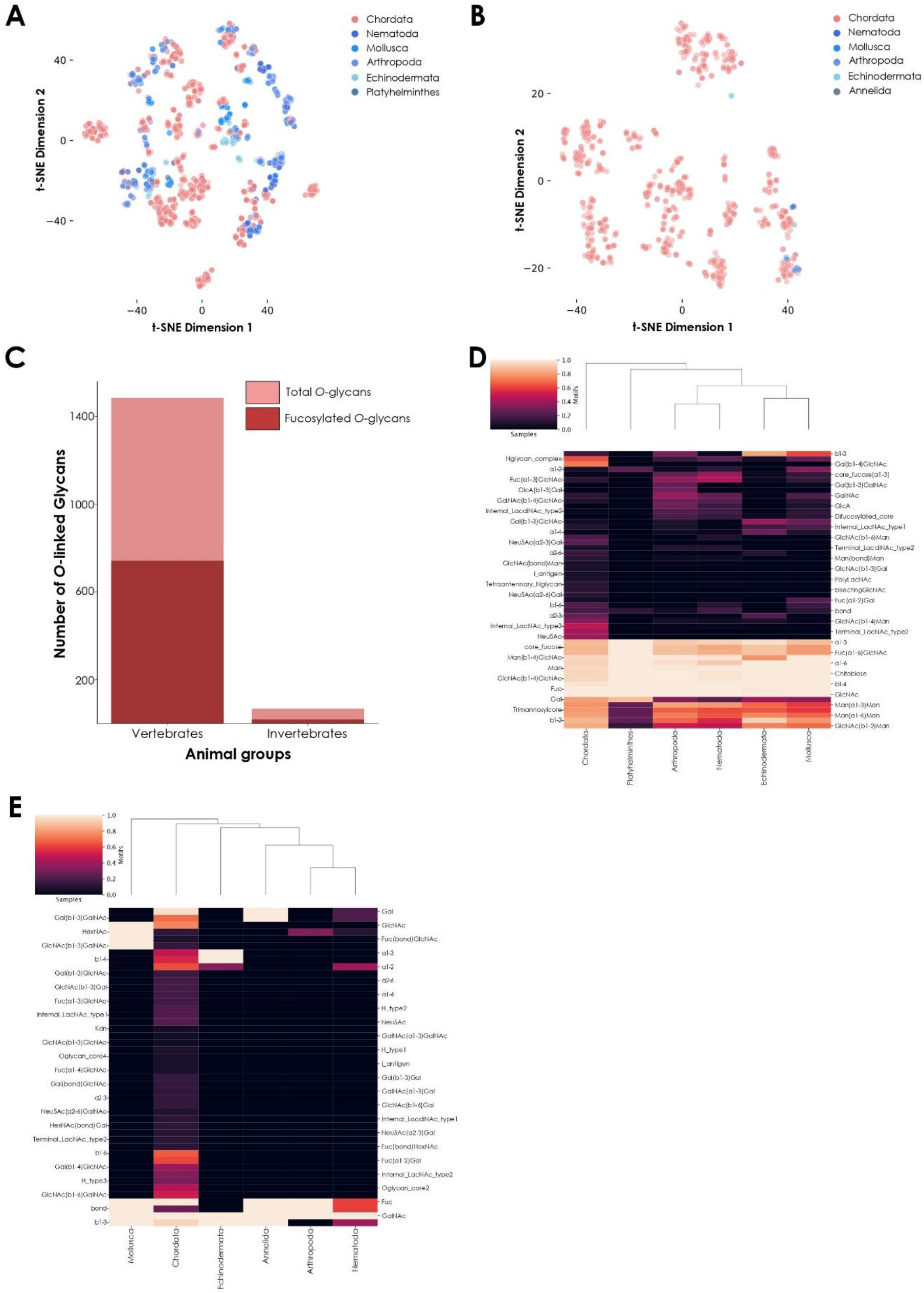
Comparison of fucosylated *N*-linked and *O*-linked glycans from vertebrates and invertebrates. **A)** Fucosylated *N*-linked glycans from animals. Glycans are represented via t-SNE as colored dots, based on their learned similarity from a SweetNet model. Colors indicate the phylum from which they stem. **B)** Fucosylated *O*-linked glycans from animals. Similar to (A), glycans are visualized via their learned similarities and colored according to their phyla. **C)** Number of total versus fucosylated *O*-linked glycans. Bar graph comparing the number of total versus fucosylated *O*-glycans from vertebrate and invertebrate species. **D)** Heatmap of fucosylated *N*-linked glycans from animals. This heatmap represents the relative abundance of different motifs from fucosylated vertebrate and invertebrate *N*-glycans. **E)** Heatmap of fucosylated *O*-linked glycans from animals. Similar to (D), the relative abundance of different motifs from fucosylated vertebrate and invertebrate *O*-glycans is shown on a heatmap.

Invertebrate *N-*linked fucosylated glycans formed distinct clusters (Fig. 3A) while their few *O-*linked fucosylated glycans were mostly colocalized with Chordata glycans, suggesting a stronger similarity on the level of (known) *O*-linked glycans (Fig. 3B). From this observation, we could deduce that most of the differences between currently known fucosylated glycans from vertebrates and invertebrates arise from *N*-linked rather than *O*-linked glycans. However, the high number of known *N*-linked glycans in both animal subgroups also resulted in some overlapping of clusters. These observations suggested at least some degree of conservation of core motifs between vertebrates and invertebrates. Using a heatmap, we first showed that many motifs were indeed conserved across all animals (Fig. 3D). These motifs are known to be specific to *N*-glycans, as exemplified by the extended chitobiose motif Man(ß1-4)GlcNAc(ß1-4)GlcNAc. We also observed that shared fucosylated glycans between vertebrates and invertebrates often contained the classical Fuc(α1-6) motif commonly found in the animal *N-*linked glycan core. Yet we also noticed some specific motifs, such as Fuc(α1-2), which in part caused the independent clustering of Mollusca. Even among invertebrate organisms, mollusks were the most prone to connect fucose using this type of linkage, resulting in distinct clustering. Another particularity of invertebrate species shown by the heatmap was the specific usage of a difucosylated core, a concomitant usage of α1-6 and α1-3 core fucosylation, in Nematoda, Arthropoda, and Mollusca, a motif usually not observed in vertebrates.

We also noticed that some invertebrate *O*-linked glycans were not completely overlapping those from vertebrates, justifying their closer examination. As only nineteen fucosylated *O*-linked glycans were identified within invertebrates in our dataset, we manually investigated their architecture and composition, in addition to representing them on a heatmap (Fig. 3E). Interestingly, from all animals in our dataset, only glycans from the nematode species *Toxocara canis* and *Toxocara cati* contained the methylated fucose variant Fuc2Me (also referred as 2-*O*-Me-Fuc), even when considering both *O*-linked and *N*-linked glycans. Little is known about this modified monosaccharide, except that it is part of highly antigenic motifs (Khoo et al., 1991). Notably, we further identified a report of the presence of this variant in the model organism *Caenorhabditis elegans* (Gue et al., 2001), reinforcing its usage within the Nematoda phylum.

These methylated monosaccharides have also been detected in another invertebrate, the annelid *Alvinella pompejana* (Talmont and Fournet, 1991). This organism lives close to deep-sea hydrothermal vents and integrates mono- and polymethylated fucose moieties into the carbohydrate part of their secreted tubes. Unfortunately, the structures of these glycans remain unsolved so far, which is why they were not part of our analysis. Of special interest are animals from the Echinodermata phylum, considered to be at the intersection between deuterostomes and protostomes, a privileged place in evolution to help us understand the differences between higher and lower animals. Interestingly, a recent publication about the echinoderm *Ophiactis savignyi* (brittle star) highlighted the presence of typical features from both vertebrate and invertebrate glycans (Eckmair et al., 2020). Monosaccharides (including fucose) from this and other echinoderms are sulfated (Vasconcelos and Pomin, 2017). The presence of fucosylated chondroitin sulfate in sea cucumber species is noteworthy. These glycosaminoglycans contain mostly 4-sulfated and 2,4-disulfated fucose residues in *Pearsonothuria graeffei* and *Isostichopus badionotus*, respectively (Chen et al., 2011), showcasing the diversity of fucose variants.

### Fucose usage across bacterial species from the microbiota

Similar to animals, bacteria also synthetize fucosylated polysaccharides. One of the most studied examples is *Helicobacter pylori*, a bacterium established in the stomach of its human host known to, occasionally, infiltrate the mucus layer and cause clinical symptoms. This bacterium is well-known for producing fucosylated glycans, mimicking Lewis blood group antigens that are also present at the surface of the stomach epithelium (Altman et al., 2005; Matos et al., 2021). The expression of host-like glycans by this organism is believed to be involved in immune evasion and one can expect that the glycan profile of many animal-associated bacteria should be adapted to this aim. To determine how fucose is used by different bacterial species, we compared the proportion of fucosylated glycans from four well-studied species known to colonize different human organs: *Helicobacter pylori*, *Pseudomonas aeruginosa*, *Staphylococcus aureus*, and *Escherichia coli* (Table 1). Our results showed that the computed ratio of fucose-containing versus total glycans can vary widely, even between species colonizing the same host (from 18.7 to 61.7%). One explanation could be an adaptation of fucose usage to the glycan profile of the organs where these organisms grow.

**Table 1.** Proportion of fucose-containing glycans in well-known human-associated bacteria. Ratios of fucosylated versus total glycans were computed and represented for four well-studied bacterial species colonizing human organs: *H. pylori*, *P. aeruginosa*, *S. aureus*, and *E. coli*. Species are sorted by decreasing proportion of fucose-containing glycans and preferred organs in human hosts are indicated.

| Bacterial species | Number of glycans | Number of fucosylated glycans | Ratio (%) | Preferred localization |
| --- | --- | --- | --- | --- |
| *Helicobacter pylori* | 94 | 58 | 61.7 | Stomach |
| *Pseudomonas aeruginosa* | 291 | 109 | 37.5 | Airways |
| *Staphylococcus aureus* | 28 | 9 | 32.1 | Skin, nasal cavity |
| *Escherichia coli* | 1,049 | 196 | 18.7 | Lower intestine |

We then decided to conduct the same analysis on all 729 bacterial species present in our dataset. To ensure sufficient representativeness and avoid biases during the calculation of ratios, we filtered species based on the number of associated glycans, keeping only those for which we had at least 30 available glycans. This selection resulted in the identification of 55 species that inserted fucose in 0-70% of their glycans. In addition, for each species, we also indicated their preferred growing environments and their known involvement in animal or plant pathologies (Supplementary Table 1). From this list, only seven species inserted fucose into more than half of their glycans and were categorized as “high-fucose content bacteria”: *Pseudomonas fluorescens* (70% of fucosylated glycans), *Rhizobium etli* (67.6%), *Helicobacter pylori* (61.7%), *Sinorhizobium fredii* (54.8%), *Bradyrhizobium japonicum* (54%), *Pectinatus frisingensis* (51.5%), and *Sinorhizobium sp* (51%). With the notable exception of *H. pylori* living in the stomach of its host and *P. frisingensis* found in beer spoilage, we noticed that all these bacteria are known to be associated with plant roots. In contrast, 34 species were categorized as “low-fucose content bacteria” (0.5-50%). From these species, only *Azospirillum brasilense* (36.8%) and *Rhizobium leguminosarum* (15.4%) are growing in association with plant roots. Most of the other organisms are found in a diversity of environments, ranging from animal or plant hosts to water, soil, or food. In this category, we also found well-known human pathogens able to infiltrate the respiratory, urinary, or gastrointestinal tracts. Some examples include *Vibrio cholerae*, *Yersinia pestis*, *Shigella dysenteriae*, or *Pseudomonas aeruginosa*.

Finally, the last fourteen species studied here were completely devoid of fucosylated glycans and their large majority is, again, known to be responsible for animal diseases. It is interesting to note that only three of these (*Lactococcus lactis*, *Leuconostoc mesenteroides*, and *Bacillus subtilis*) grow in association with plants and only *B. subtilis* is known to be associated with their roots, in addition to infecting animals. *L. lactis* is used for industrial activities but was initially found on plant leaves and *L*. *mesenteroides* is found on fruits and vegetable skin that it can colonize. Comparing root-associated bacteria with the rest, we found a clear tendency of this group to exhibit fucosylated glycans compared to other bacterial species (Fig. 4A). Taken together, these results indicated a strong correlation between the ratio of fucose-containing glycans from different bacterial species and their preferred growing environment as well as pathogenic potential.

**Figure 4.**
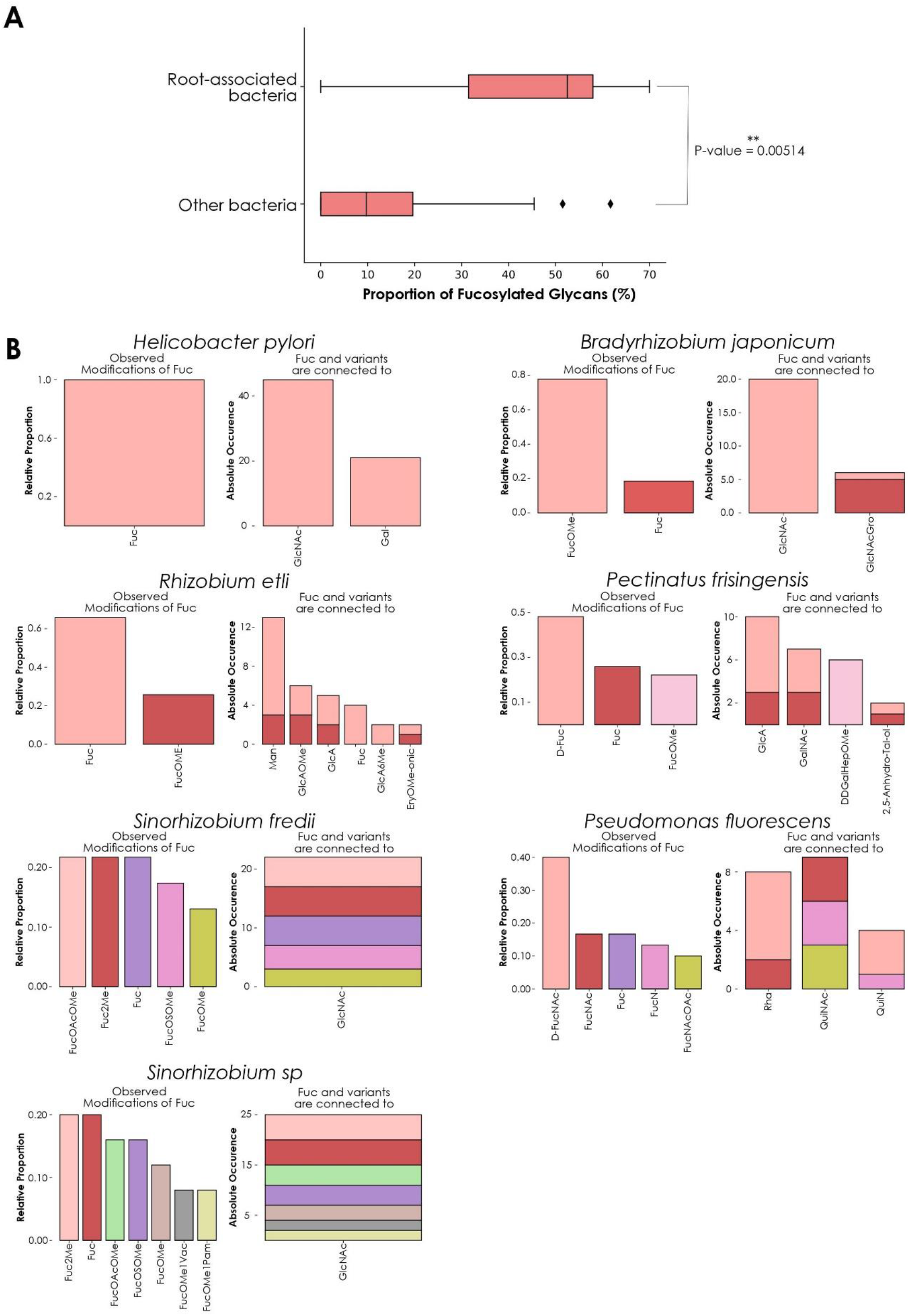
Fucose distribution and usage in glycans among high-fucose content bacterial species. **A)** Comparing fucose content across bacterial glycans. This boxplot represents the percentage of fucose-containing glycans from root-associated bacterial species (top) versus all other bacteria (bottom). Only bacteria that contain at least 30 glycans in our dataset are displayed. The displayed p-value was computed with a one-tailed Welch’s t-test. **B)** Preferred neighbors of fucose in glycans from high-fucose content bacteria. Each graph represents the relative abundance of fucose and fucose variants. Colored stacked bars correspond to the occurrence of the preferred neighbors of fucose variants in glycans, with colors corresponding to the variants observed on the left part of each graph.

As *H. pylori* appeared as an exception to this trend, we compared its fucosylated glycans to the other high-fucose content bacteria by analyzing preferred fucose neighbors and linkages within its glycans (Fig. 4B). Fucose usage and preferred connections observed in *H. pylori* glycans mirrored the characteristic fucosylation features from its human host. While other bacteria within this group use at least one fucose variant, *H*. *pylori* only inserts classical L-fucose residues into its glycans. We also noticed that in this organism, fucose is connected to GlcNAc and Gal in proportions mimicking its usage observed in animals (Fig. 1D), consistent with the known role of glycans from this species in mimicking and evading the human immune system.

In contrast, *P. frisingensis* and the root-associated bacteria displayed a large variety of fucose variants (including D-Fuc, D-FucNAc, FucNAc, FucN, FucNAcOAc, as well as the methylated forms FucOMe, FucOAcOMe, Fuc2Me, FucOMe1Vac, FucOMe1Pam, and FucOSOMe) that they connect to an even larger collection of modified monosaccharides. Intriguingly, *P. fluorescens* connects fucose variants to monosaccharides that were absent from the other species: Rha, QuiNAc, and QuiN. These observations demonstrated that, even if all these species are associated with plant roots (with the notable exceptions of *H. pylori* and *P. frisingensis*), the way they construct their fucosylated glycans can differ. Such behavior can be explained by the requirement of adapted glycans interacting with the roots of different plant or fungal species also colonizing the same soil environments. It is interesting to note that methylated fucose is often used as an inhibitor of fucose-binding lectins because of stronger interactions with these proteins than fucose (Chemani et al., 2009). This is particularly meaningful as we know that most fucose-binding lectins used for experimental purposes come from plant or fungal species. Taken together, this could point to a role of methylated fucose in a specific recognition system required to establish bacteria-plant symbioses.

### Role of fucosylated glycans in the pathogenic potential of *E. coli* strains

Similar to *H. pylori, Escherichia coli* is a well-studied human-associated bacterium usually found growing in the lower intestine as part of the normal microbiota. It is known that under certain circumstances, *E. coli* can infiltrate different tissues, resulting in infections and diseases of the gastrointestinal and urinary tracts. Further, certain *E. coli* strains are more capable of inducing a pathologic situation than others (for a review, see Riley, 2020). Given the importance of fucosylation in escaping the immune system and the correlation between the amount of fucosylated glycans and the previously highlighted habitat specificities of bacteria, we decided to investigate the role of fucosylated glycans on the pathogenic potential of *E. coli* strains.

For this, we started with a set of strains annotated as pathogenic or non-pathogenic. Representations of total (Fig. 5A) and fucosylated (Fig. 5B) glycans from pathogenic and non-pathogenic bacteria showed that, while total glycans from both categories were widely dispersed and often overlapped, the representation generated using only fucosylated glycans showed a clearer segregation of the dots according to the known pathogenic capacities of the strains they stemmed from.

**Figure 5.**
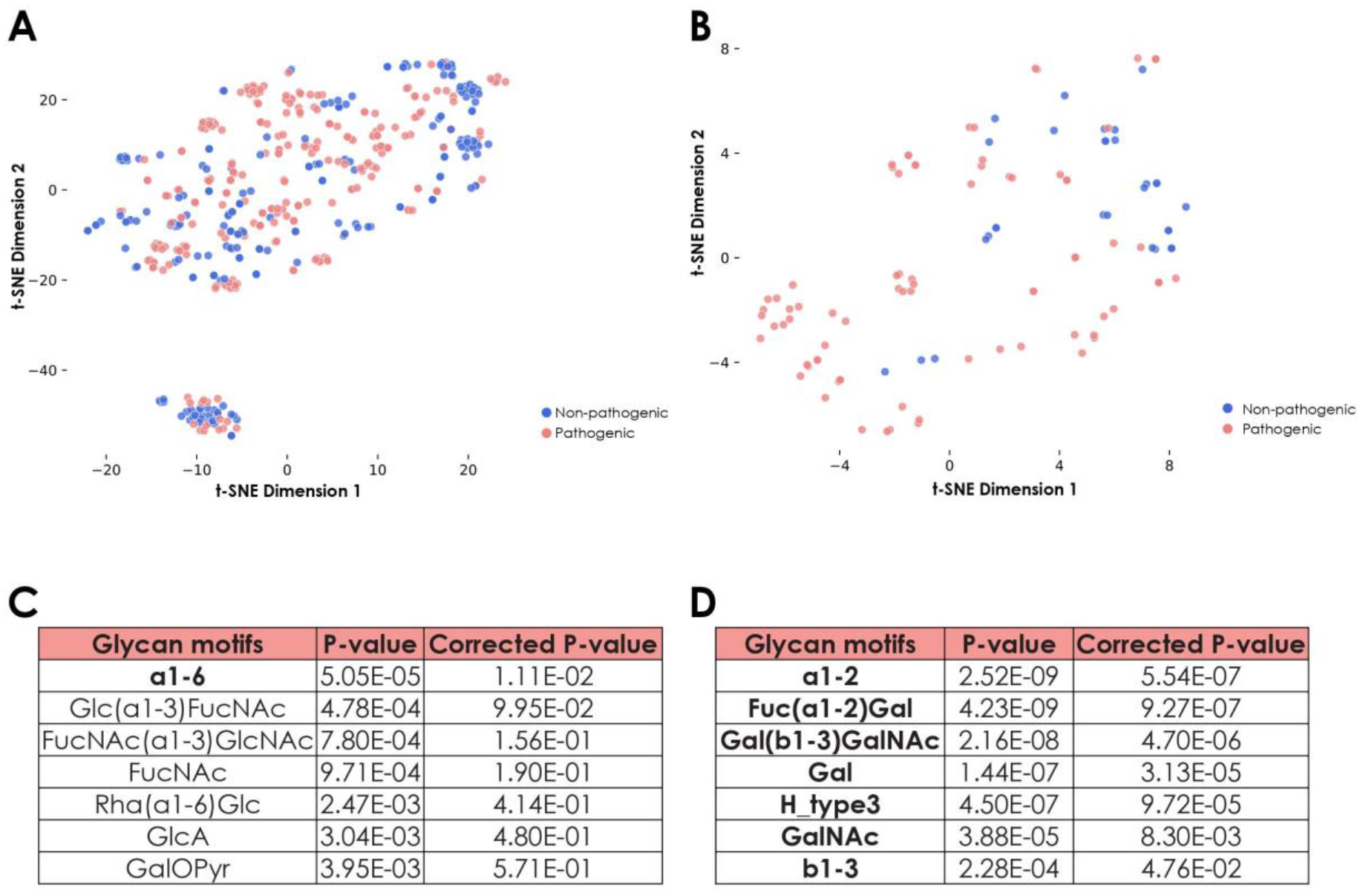
Specific features of fucosylated glycans from pathogenic and non-pathogenic *E. coli* strains. **A)** Representation of all glycans from pathogenic and non-pathogenic *E. coli* strains. Shown is a t-SNE graph representing all glycans from both pathogenic and non-pathogenic *E. coli* strains via their learned similarities by a SweetNet model. Colors indicate the group from which they stem. **B)** Representation of fucosylated glycans from pathogenic and non-pathogenic *E. coli* strains. Fucose-containing glycans from both groups of *E. coli* strains are visualized with their learned similarity via t-SNE. **C)** Identification of specific motifs from fucosylated glycans to distinguish groups of *E. coli* strains. To determine specific motifs from fucose-containing glycans for non-pathogenic versus pathogenic *E. coli* strains, we computed their statistical enrichment with one-tailed Welch’s t-tests and a Holm-Šídák correction for multiple testing. Motifs indicated in bold have a corrected p-value inferior to 0.05. **D)** Identification of specific motifs for pathogenic *E. coli* strains. Similar as in (C), we computed statistical enrichment of motifs from pathogenic versus non-pathogenic *E. coli* strains.

As this distribution suggested that composition and architecture of fucosylated glycans are tightly associated with the pathogenic capacities of the corresponding strains, we further investigated these features by characterizing enriched motifs (Fig. 5C-D). The most striking differences pointed to the preferred usage of FucNAc as well as the FucNAc(α1-3)GlcNAc and Glc(α1-3)FucNAc motifs within the non-pathogenic strains (Fig. 5C). Conversely, *E. coli* strains that are more likely to trigger an infection were also more prone to display Fuc(α1-2) Gal motifs (Fig. 5D). Pathogenic strains minimized the usage of FucNAc, which is not found in humans, while they favored the addition of terminal Fuc(α1-2)Gal, mimicking motifs observed in human Lewis antigens. This presumably serves to evade the immune system during infection. Similarly, the Forssman antigen that contains the trisaccharide GalNAc(α1-3) GalNAc(ß1-3)Gal, and which is present in different mammalian species including some humans (Yamamoto et al., 2012), was also exclusively found in glycans from pathogenic strains, reinforcing the essentiality of the mimicking process to confer pathogenicity.

### Fucosylated glycans from plants

Despite the lower number of plant glycans in our database (2,873), compared to animals (8,260) and bacteria (8,072), we applied similar methods to determine how members of this kingdom integrate fucose in their glycans. Via the order-level heatmap plus complementary manual investigations, fucose usage appeared relatively conserved across plant species (Fig. 6A). Most species use the classical L-fucose and insert it into their usual core fucose Fuc(α1-3)GlcNAc motif. The absence of chitobiose, as the common core of *N*-linked glycans, from eight plant orders in our dataset (Santalales, Nymphaeales, Gleicheniales, Asterales, Ipomoea, Gentianales, Selaginellales, and Equisetales) indicated the absence of recorded fucosylated *N*-linked glycans from these organisms in our dataset. This suggests the need for more study in this taxonomic subset of the Plantae kingdom. We also observed differences in the Gentianales and Ipomoea taxa, which prefer to use specific fucose variants. Species from the former group are often toxic lianas that insert D-FucOMe within their glycans and are the only ones from our database that also incorporate the D-FucOAcOMe variant (only observed in *Periploca sepium*).Organisms from the Ipomoea order rather insert D-Fuc, especially within the Rha(α1-2)D-Fuc motif, a behavior also observed among members of the Asterales order.

**Figure 6.**
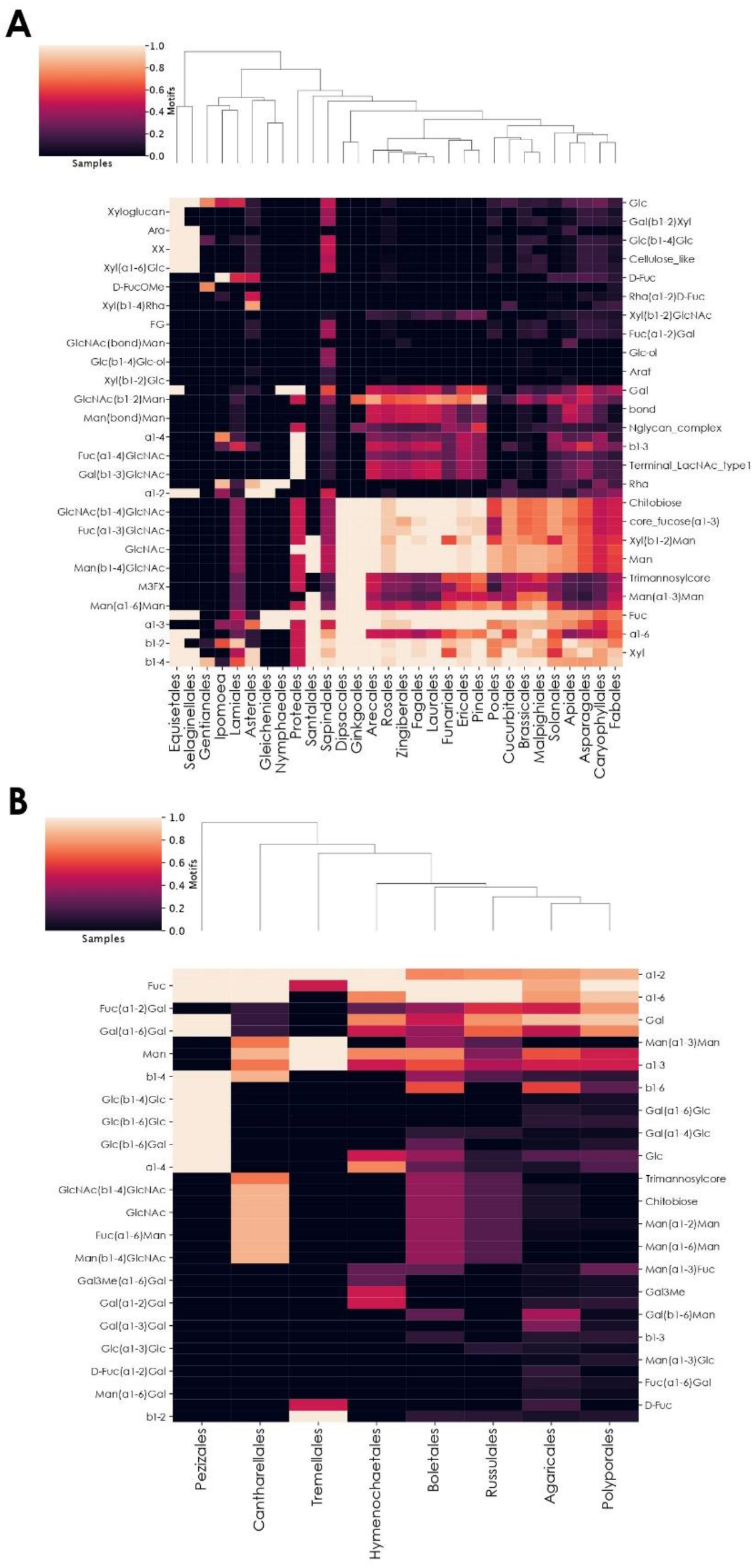
Preferential motif usage in fucosylated glycans across plant and fungi orders. **A-B)** Heatmaps of motifs in fucosylated glycans at the plant order-level (A) or fungi order-level (B). These heatmaps represent the relative abundance of different motifs from fucosylated plant (A) or fungal (B) glycans in different taxonomic orders.

### Fucosylated glycans from fungi

Among the seven investigated kingdoms, Fungi contained the second-lowest proportion of fucosylated glycans (Fig. 1A). These organisms differed from animals, plants, and bacteria by preferentially connecting their fucose residues to galactose via α1-2 linkages. However, through an order-level heatmap (Fig. 6B), we noticed that two groups seemed to be devoid of this characteristic motif in our dataset: Pezizales and Tremellales. A closer manual inspection revealed that only one glycan from Pezizales was fucosylated in our database, mirroring the situation observed with the Annelida phylum. Similarly, only two of the 184 glycans available for the Tremellales order were fucosylated. Compared to other kingdoms, Fungi organisms contain fucose within specific motifs, including Fuc(α1-6)Man, Man(α1-3)Fuc, D-Fuc(α1-2)Gal, as well as Fuc(α1-6)Gal. Similar to plants, the chitobiose motif was restricted to Cantharellales, Boletales, Russulales, and, in lower proportions, Agaricales. This indicated the absence of fucosylated *N*-linked glycans from the four other orders in our dataset. Finally, it is important to note that only eight of the 58 Fungi orders in our dataset were represented on this heatmap at all, as the other 50 entirely lacked fucosylated glycans. This is consistent with the low proportion of fucosylated glycans among fungi species, including the known lack of fucosylated glycans in the model organism *Saccharomyces cerevisiae*, a member of the non-represented Saccharomycetales order (Chigira et al., 2008).

### Role of fucose-associated enzymes in human diseases

Differences in the usage of fucose, which, as shown, is often connected to disease-relevant contexts, stem from differences in the biosynthesis of glycans. FUT (fucosyltransferase) genes encode glycosyltransferases that specifically transfer fucose monosaccharides onto glycans. Thirteen of these genes are present in humans and the encoded enzymes insert fucose residues into elongating glycans through different linkages. As they are known to be often associated with diseases (Schneider et al., 2017), we decided to extract available experiments and their differential expression data from the EMBL-EBI Expression Atlas concerning these genes (https://www.ebi.ac.uk/gxa/home, Papatheodorou et al., 2018).

To investigate whether and how fucosyltransferases are impacted during embryonic development, disease progression, environmental stress response, or other conditions, we searched the 3,903 datasets on Expression Atlas database for the 758 experimental conditions in which at least one fucosyltransferases was up- or down-regulated (Supplementary Table 2). FUT8 was the most frequently identified gene (in 251 experiments), while FUT5 was particularly underrepresented (in eight experiments, Fig. 7A). As FUT8 was previously identified in many different cancer-related studies (Huang et al., 2021; Ma et al., 2021; Bastian et al., 2021), we investigated in which proportion fucosyltransferases as a whole were involved in cancer. From the eight experiments in which at least nine FUT genes were differentially expressed together, six were cancer studies. Compared to the 229 datasets with “cancer” in their title on the EMBL-EBI website, representing 5.9% of the whole database, we identified 12% in our list of 758 FUT-related studies. Interestingly, this value increased from 12 to 13.6% when we considered only the 369 experiments in which at least two fucosyltransferases were differentially expressed. These results indicated an enrichment of cancer-related studies that were associated with fucosyltransferases, compared to the whole database. It is important to note that these computed proportions of cancer-related experiments are an underestimate, as we restricted this analysis to titles explicitly mentioning “cancer”. Many cancer studies rather indicate the cancer type, such as “adenocarcinoma”, “sarcoma”, or “leukemia”, and were consequently not considered in these calculations.

**Figure 7.**
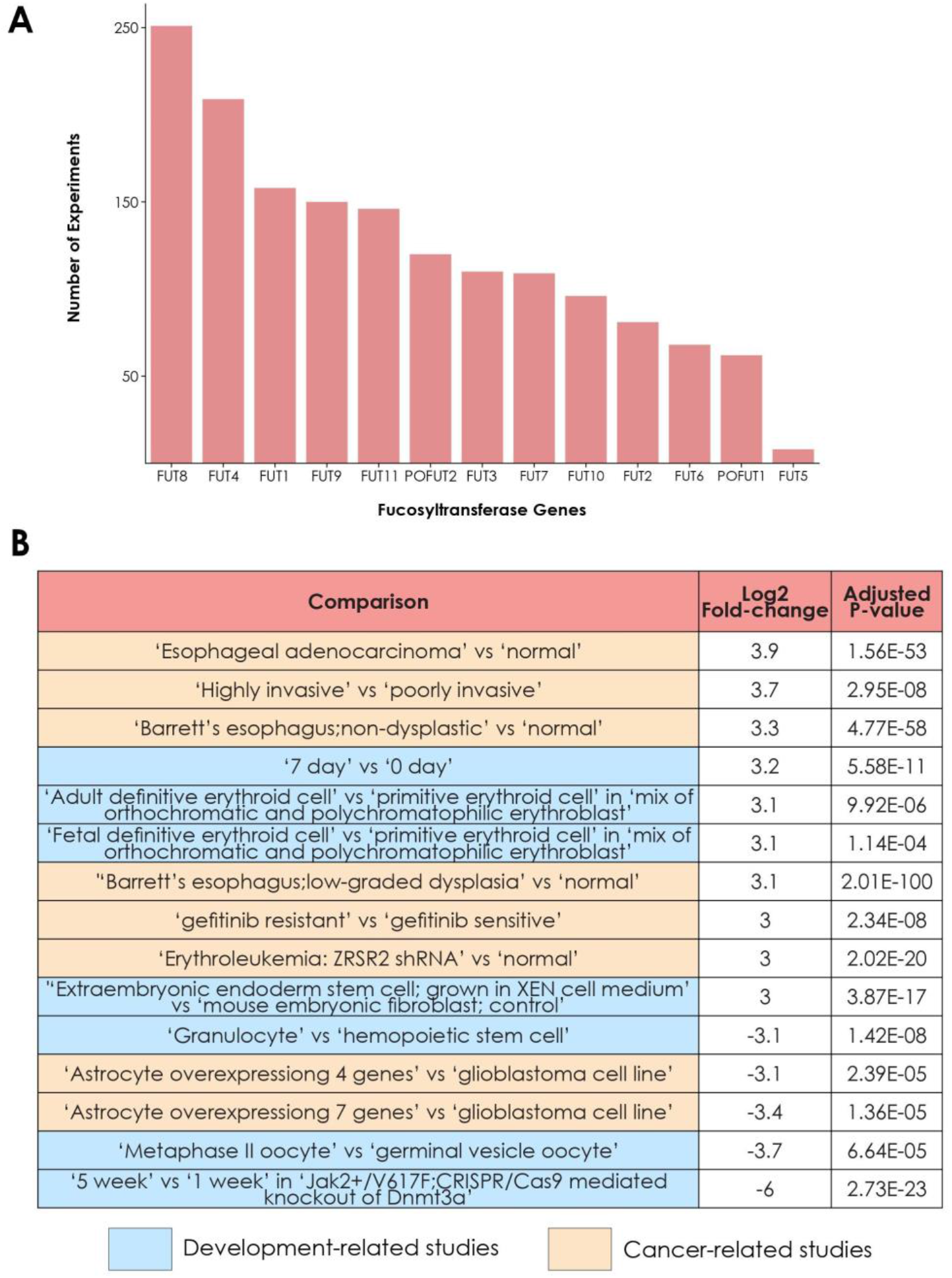
Involvement of fucosyltransferases in different biological processes and diseases. **A)** Frequency of differential expression of fucosyltransferases in different experiments. The number of experiments in which each of the thirteen human fucosyltransferases are differentially expressed is represented as a bar graph. **B)** Experiments in which FUT8 is highly differentially expressed. This table contains the comparisons from experiments in which FUT8 is differentially expressed with an adjusted p-value inferior to 0.05 and a log2(fold-change) either superior or equal to 3 or inferior or equal to −3. Comparisons highlighted in blue came from development-related studies. Comparisons highlighted in yellow came from cancer-related studies.

Knowing the particular importance of FUT8, encoding the enzyme responsible for core fucosylation of *N*-linked glycans, we further manually investigated the 251 experiments in which FUT8 was differentially expressed. We found a total of at least 98 experiments either mentioning cancer or studying particular cancers or cancerous cell-types. Other frequent experiments studied development, infections, as well as studies of immune processes and immune-related diseases. In addition, we noticed that FUT8 was strongly differentially expressed (log2 fold-change either superior or equal to 3, or inferior or equal to −3) in fifteen comparisons from eleven experiments (Fig. 7B). Eight of these comparisons were extracted from cancer-related studies, and the others originated from studies on stem cells or differentiation. Interestingly, in the eight cancer studies, FUT8 was always upregulated in the pathologic condition versus control. Taken together, these results point to a strong association between fucose metabolism and human pathologies, especially cancer. It also confirmed that animal hosts are responding to infection by modulating fucose-related pathways, validating that this monosaccharide is important for both the effector and the target of the infection.

## DISCUSSION

In this work, we demonstrated that, despite its status as a ubiquitous monosaccharide conserved across all kingdoms, fucose can be modified, inserted, and used in many different ways. We presented two major trends in the way fucose is used in organisms. The first one was observed in commensal species that mimic host glycans to be recognized as a host “self” entity. This is the case of the two bacterial species *Helicobacter pylori* and *Escherichia coli*, which are both colonizers of humans and display glycan structures mimicking Lewis blood group antigens to avoid the immune system. There is also another trend, leading some organisms to develop and use highly specific fucosylated glycans through the elaboration of original variants (i.e., methylated fucose) or the insertion of fucose within uncommon motifs. This could be a way for symbiotic organisms to specifically recognize their partners or to attract species that evolved highly specific recognition systems.

### Limitations of studying taxonomic fucose distribution regarding data availability

It is important to note that the comparative analysis of monosaccharide usage across many different organisms from distant kingdoms is limited by the level of data availability. In this study, we repeatedly noted that, for some organisms, the amount of available data is insufficient to make any insightful conclusion. This was illustrated by the Annelida phylum which contained, in our database, only one fucosylated glycan. We know that additional fucose-containing polysaccharides exist in this group, as demonstrated by the study of *Alvinella pompejana* (Talmont and Fournet, 1991). Yet, because glycan structures were not available, they could not be considered here. Consequently, it is important to remember that, while the presence of motifs constitutes a signal of high confidence, absence of motifs or designating a motif as “specific” is ultimately dependent on an assumption of reasonably complete data. A motif can be absent in a species because it does not exist or because it simply has never been observed so far. This can lead to important biases mirroring our level of knowledge about different species. Some taxa are not well, or even not at all, represented in our dataset, strictly reflecting the academic literature. As a consequence, this missing critical mass of information raises many uncertainties. This situation was well exemplified here by the gap in our knowledge when comparing well-studied animals and bacteria versus less well-studied plants, fungi, protista, and archaea. For this reason, it will be important for future analyses to have access, for each species, to a representative profile of its glycans and, for each phylum, to a panoply of representative species. Fortunately, with continuous efforts spent towards analyzing glycans in an ever-increasing number of species, this limitation is perpetually being pushed back.

### Differential usage of *O*-linked glycans in invertebrate species

In vertebrates, fucosylated *O*-linked glycans are present on many proteins, notably secreted proteins from the mucin family. Our analyses showed that invertebrates seemed to exhibit a low abundance of fucosylated *O*-linked glycans in our dataset. However, proteins such as mucins also exist in invertebrates and seem to carry different *O*-linked glycans than those in vertebrates (Staudacher, 2015), suggesting the need for more research in this area. In vertebrates, *O*-linked glycans are important for the immune system and for the synthesis of mucus (Giovannone et al., 2018; Nason et al., 2021), a substance modulating pathogen adherence in animals (Gonzalez-Morelo et al., 2020). As a consequence, disruption of this type of glycosylation is associated with many human diseases (Hansson, 2019).

From our observations, provided this is not a consequence of incomplete data, the question that remains is to determine how invertebrates deal with this apparent depletion of fucosylated *O*-glycans. Also, if fucosylated *O*-glycans are really depleted in invertebrates, we can reasonably hypothesize that this should be reflected by the presence/absence of the corresponding enzymes or at least by variations in their expression level compared to vertebrates. These statements remain to be assessed in future work to further investigate this hypothesis.

### Role of fucose in the association of bacteria to plant roots

A priori, one could expect organisms that colonize similar environments to have similar requirements. Accordingly, we for example showed that *H. pylori* and pathogenic *E. coli* strains both mimicked Lewis blood group antigens from their human host to avoid the immune response. In the case of root-associated bacteria, their divergent set of fucosylated glycans, that also did not mirror the preferred fucose usage in plants, seems to exclude a mimicking process. One explanation is that plants display an altered immune response compared to animals that, potentially, alters the incentives and mechanisms of molecular mimicry. On the other hand, the root-associated bacteria analyzed here are symbiotic organisms from the rhizobiome, rather than pathogens, and might not require mimicry.

Despite the large variety of fucosylated glycans produced by root-associated bacterial species, we noticed a common usage of methylated fucose variants. Together with the fact that most of the fucose-binding lectins are extracted from plants and fungi, we propose that root-associated bacteria express fucosylated glycans to be recognized by plant lectins. In addition, differences observed in bacterial species could be related to their association with plants from very distinct taxa that express different glycan profiles. As glycan profiles can be different between organs in animals, an analogous phenomenon in plants could also help explain why high-fucose content bacteria are specifically suited to interact with roots, while low-fucose content bacteria (except *Bacillus subtilis*) seem to be more adapted to interact with higher parts of the plant, such as leaves or fruits. This could be an interesting parallel with the observation that the high-fucose content bacterium *H. pylori* can infect the human stomach while low-fucose content bacteria rather colonize respiratory, urinary, and gastrointestinal tracts.

During our investigation of the specificities observed in fucosylated *O*-linked glycans from invertebrates, we noticed the specific usage of methylated fucose, Fuc2Me, in several nematode species (*Toxocara canis*, *Toxocara cati*, and *Caenorhabditis elegans*). It is interesting to note that many nematode species share the soil environment and may also rely on the use of methylated fucose to survive in such a habitat. *Meloidogyne incognita*, for instance, is a soil-resident root-knot nematode known to parasitize plant species. Efforts to determine how these animals detect their host have demonstrated that they are attracted by secreted plant glycans (Tsai et al., 2021). Future work on this nematode and other soil-resident organisms should be helpful to determine if they also use Fuc2Me to grow in the soil and to which extent their own glycans are important to establish a parasitic relation with their plant host. For now, we already know that some organisms can produce lectins specifically adapted to target methylated glycans from other species (Wohlschlager et al., 2014). This mechanism serves as an innate defense system and underlines the complex relations that can occur between resident organisms from a shared environment.

### Predicting bacterial strain pathogenicity based on fucose usage

Based on the observations made with root-associated bacteria, we showed that fucose usage was also correlated with the preferred growing environment of bacteria and their capacity to trigger an infection. We analyzed data from several *E. coli* strains annotated as pathogenic or non-pathogenic to identify differences concerning their fucose usage and showed that pathogenic bacteria were more likely to produce glycans similar to their host. This is known as a mimicking mechanism used by microorganisms to escape the host immune system (Carlin et al., 2009). Consequently, we showed that it was possible to separate pathogenic from non-pathogenic *E. coli* strains solely based on their fucose-containing glycans. Including all known glycans did not improve the segregation of bacteria according to their pathogenic potential. This observation suggested that, at the glycan level, most of the peculiarities of pathogenic versus non-pathogenic strains seem to rely on fucose-containing glycans. This was for instance confirmed by the tendency of pathogenic strains to mimic Lewis blood group antigens. Such a result indicates that it could be promising to train a machine-learning model predicting the pathogenic potential of bacterial strains using only their fucosylated glycans, complementing other results such as the already identified rhamnose enriched in non-pathogenic strains (Bojar et al., 2021).

## DATA AVAILABILITY STATEMENT

Data used in this study can be freely accessed at https://github.com/BojarLab/glycowork and in the supplementary tables.

## AUTHOR CONTRIBUTIONS

Conceptualization: D.B., Data Curation: L.T., D.B., Funding Acquisition: D.B., Investigation: L.T., D.B., Resources: D.B., Software: L.T., D.B., Supervision: D.B., Visualization: L.T., D.B., Writing—Original Draft Preparation: L.T., D.B., Writing—Review & Editing: L.T., D.B.

## FUNDING

Branco Weiss Fellowship – Society in Science awarded to D.B.; Knut and Alice Wallenberg Foundation; University of Gothenburg, Sweden

## ACKNOWLEDGMENTS

The authors would like to thank Jon Lundstrøm and Emma Korhonen for helpful feedback and comments.

---

**Supplementary Table 1. Relations between the pathogenic potential, growing environment, and proportion of fucose-containing glycans from bacterial species.** The 55 bacteria, filtered based on the number of known glycan structures, are ordered by increasing ratio of fucosylated versus total glycans. For each bacterium, preferred localization, associated disease, and corresponding references are indicated.

**Supplementary Table 2. List of experiments from the EMBL-EBI Expression Atlas in which at least one FUT gene is differentially expressed.** The total 758 experiments in which at least one of the thirteen FUT genes is differentially expressed are ordered by increasing number of significantly up- or down-regulated FUT genes. For each experiment, the list of differentially expressed FUT genes is indicated.

## Notes

https://github.com/BojarLab/glycowork

### Competing Interest Statement

The authors have declared no competing interest.

